# Deciphering the genetic links between NAFLD and co-occurring conditions using a liver gene regulatory network

**DOI:** 10.1101/2021.12.08.471841

**Authors:** Sreemol Gokuladhas, William Schierding, Tayaza Fadason, Murim Choi, Justin M. O’Sullivan

**Author notes:** Corresponding author: Justin Martin O’Sullivan.

## Abstract

**Background & Aims:** Non-alcoholic fatty liver disease (NAFLD) is a multi-system metabolic disease that co-occurs with various hepatic and extra-hepatic diseases. The phenotypic manifestation of NAFLD is primarily observed in the liver. Therefore, identifying liver-specific gene regulatory interactions between variants associated with NAFLD and multimorbid conditions may help to improve our understanding of underlying shared aetiology.

**Methods:** Here, we constructed a liver-specific gene regulatory network (LGRN) consisting of genome-wide spatially constrained expression quantitative trait loci (eQTLs) and their target genes. The LGRN was used to identify regulatory interactions involving NAFLD-associated genetic modifiers and their inter-relationships to other complex traits.

**Results and Conclusions:** We demonstrate that *MBOAT7* and *IL32*, which are associated with NAFLD progression, are regulated by spatially constrained eQTLs that are enriched for an association with liver enzyme levels. *MBOAT7* transcript levels are also linked to eQTLs associated with cirrhosis, and other traits that commonly co-occur with NAFLD. In addition, genes that encode interacting partners of NAFLD-candidate genes within the liver-specific protein-protein interaction network were affected by eQTLs enriched for phenotypes relevant to NAFLD (*e*.*g*. IgG glycosylation patterns, OSA). Furthermore, we identified distinct gene regulatory networks formed by the NAFLD-associated eQTLs in normal versus diseased liver, consistent with the context-specificity of the eQTLs effects. Interestingly, genes targeted by NAFLD-associated eQTLs within the LGRN were also affected by eQTLs associated with NAFLD-related traits (*e*.*g*. obesity and body fat percentage). Overall, the genetic links identified between these traits expand our understanding of shared regulatory mechanisms underlying NAFLD multimorbidities.

## Introduction

Non-alcoholic fatty liver disease (NAFLD) is a chronic liver disease that has emerged as a global health burden, in parallel with obesity and type-II-diabetes(1,2). The accumulation of hepatic fat in the absence of significant alcohol consumption or infection is the defining feature of NAFLD. NAFLD includes a broad spectrum of liver pathologies ranging from simple steatosis to NASH, which predispose susceptible individuals to inflammatory fibrosis, cirrhosis, and hepatocellular carcinoma(3). Systemic metabolic disorders such as type-II diabetes, obesity, and metabolic syndrome(4–6) commonly co-occur and are widely recognized as fueling the hepatic and systemic manifestations of NAFLD.

Epidemiological studies have found that NAFLD is associated with chronic kidney disease(7,8), obstructive sleep apnea (OSA)(9,10), polycystic ovarian syndrome(11), and extrahepatic cancers(12). The bidirectional nature of the observed relationship between NAFLD and these conditions is consistent with the existence of a network of interactions where NAFLD can be both the cause and consequence of another phenotype. Candidate gene approaches, transcriptomic studies, and genome-wide association studies have provided key insights into the genetic modifiers associated with these diseases(13–16). However, there is a lack of robust evidence characterizing shared genetic risk, or the biological mechanisms through which genetic variation contributes to the observed shared etiology or multimorbidity. Despite this, the combination of the heritable nature of NAFLD and its associated diseases, and their phenotypic co-occurrence is consistent with the existence of an underlying shared genetic risk.

The regulation of gene transcription is one mechanism through which genetic variation can modify disease susceptibility and progression(17,18). Transcriptional programs are cell-type or tissue-specific and emerge through gene regulatory networks in which the transcriptional regulators (*e*.*g*., transcriptional factors, DNA regulatory elements) recruit the transcriptional machinery to genes. Chromatin architecture captures these regulatory interactions, including interactions that bring enhancer elements located elsewhere in the genome into physical proximity with promoter regions, by chromatin looping(19), inducing gene transcription(20). The chromatin interactions that are captured using high-throughput chromosome conformation capture (Hi-C)(21) include a representative snap-shot of the genome-wide network of regulatory connections occurring at the moment when the chromatin structure was captured(21). Genetic variation that disrupts this interaction (*e*.*g*. by reducing transcription factor binding affinity, or by inactivating motifs that are necessary for establishing or maintaining looping) can alter gene expression and result in an increase in an individual’s risk of developing a complex disease(s)(22,23).

We reasoned that the LGRN would include regulatory patterns that bridge NAFLD and other complex diseases. Therefore, we constructed a LGRN of eQTLs and their target genes, that was supported by chromatin contacts between the regulatory regions containing the single nucleotide polymorphisms (SNPs) and the target genes in the liver. We identified disease-specific eQTL-gene connections by determining if our findings replicated within an eQTL dataset from NAFLD patients. A network-based analysis was performed using the NAFLD eQTL integrated protein-protein interaction network (PPIN) to identify interacting partners of the NAFLD-candidate genes. Using the LGRN, we identified the total set of regulatory interactions for the NAFLD-associated eQTL target genes, NAFLD-candidate genes, and the interacting partners of the candidate genes. Our results demonstrate that the target genes of NAFLD-associated eQTLs, NAFLD-candidate genes and their interaction partners were also affected by eQTLs associated with the multimorbid traits of NAFLD. Collectively, our findings provide novel insights into the shared genetic aetiology underlying NAFLD and other complex traits with hepatic and extra-hepatic manifestations.

## Methods

### Construction of liver gene regulatory network (LGRN)

The LGRN is a network of spatially constrained regulatory interactions between eQTLs and genes in the liver. ‘Spatially constrained’ eQTLs and target genes have been captured as physically interacting in high-throughput chromatin contact datasets (Hi-C)(21). We used CoDeS3D(24), a python-based algorithm, to construct the LGRN as follows: 1) All SNPs genotyped in GTEx V8(25) (MAF>1%) were used as input to the CoDeS3D algorithm to identify physically interacting genes in three liver-specific Hi-C datasets: i) endothelial cells of the hepatic sinusoid (GEO: GSE105988)(26); ii) HepG2 (GEO: GSE105381)(26); and iii) primary liver tissue (GEO: GSE87112)(27). 2) The identified SNP-gene pairs were then queried against liver tissue genotype-expression data from the GTEx expression quantitative trait loci (eQTL) dataset (dbGaP Accession phs000424.v8.p2). This enabled the identification of eQTLs that are associated with changes in the transcript levels of the interacting target genes. CoDeS3D controls the false-positive rate for identification of eQTL-target gene associations using a Benjamini-Hochberg false discovery rate (FDR) correction. eQTL-target gene pairs with FDR<0.05 were included within the LGRN. Chromosomal locations for SNPs and genes are reported according to the human reference genome GRCh38/hg38 assembly.

### Correlation analysis of eQTLs

Pearson’s correlation analysis was used to determine if the number of eQTLs in the LGRN we identified was linearly related to the total number of common SNPs (dbSNP build 151) that were present on each chromosome of the human genome. Common SNPs were defined as any SNP with a minor allele frequency (MAF) >1% in any of the 1000 Genome project super-populations (*i*.*e*. African [AFR], Ad Mixed American [AMR], East Asian [EAS], European [EUR], South Asian [SAS])

### Identification of loss-of-function intolerant genes within the LGRN

LOEUF (loss-of-function observed/expected upper bound fraction) constraint scores were obtained from gnomAD(28) (https://gnomad.broadinstitute.org/) as an estimate of gene tolerance to mutation. LOEUF scores of < 0.35 indicate the gene has a strong intolerance to mutation, that is, loss-of-function (LoF) variants within these genes are selected against in the general population. By contrast, genes with a LOEUF score of > 0.35 are more likely to tolerate the accumulation of LoF variants.

### NAFLD-candidate gene analysis

Genes associated with NAFLD risk and severity by comprehensive transcriptome analysis, genetic risk score calculations, or replication in multiple GWAS studies were identified (Table 1), hereafter referred to as “NAFLD-candidate genes”. First, we identified the NAFLD-candidate genes that were affected by spatially constrained eQTLs within the LGRN we established. Then we performed bootstrap analysis to determine if the number of NAFLD-candidate genes identified within the LGRN was significant compared to the randomly chosen genes. Bootstrapping was performed (10000 iterations) by randomly sampling a set of genes (equivalent size to that of NAFLD-candidate genes) from a background set (*i*.*e*. all genes in the GENCODE database [version 26]). The bootstrap *p-value*, indicate the chance of finding random genes within the LGRN.

**Table 1.**
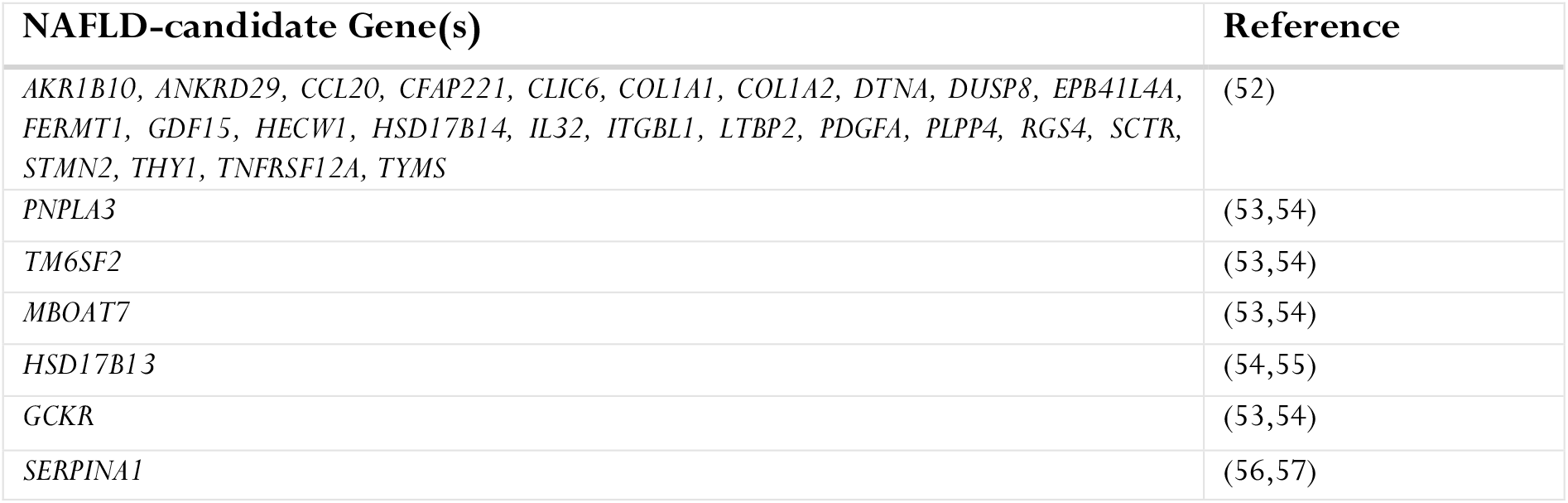
NAFLD-candidate genes previously identified by transcriptome, GWAS and polygenic risk score approaches

### Protein-protein interaction network (PPIN) analysis

The liver tissue-specific PPIN (based on GTEx RNA-seq datasets; file: https://netbio.bgu.ac.il/tissuenet2-interactomes/TissueNet2.0/GTEx-RNA-Seq.zip]) was downloaded from TissueNet v.2(29) on 05/07/2021. We then queried the liver-specific PPIN using the *ego* function in the *igraph* package(30) (version 1.2.6) to extract proteins (genes) that directly interact with the NAFLD-candidate gene encoded proteins (Supplementary data 4 table 1). The liver-specific constrained eQTLs associated with the expression change of the resulting PPIN partner protein encoding genes (*i*.*e*. partner genes) were identified using the LGRN. Subsequently, the eQTLs were queried against the GWAS catalog (downloaded on 08/07/2021). A hypergeometric test coupled with Benjamini-Hochberg FDR correction was used to identify enrichment for eQTLs associated with particular traits in the GWAS catalog.

### Identification of NAFLD-associated regulatory interactions

NAFLD-associated SNPs (N = 117) at or below the threshold suggestive of global significance (p ≤ 1 × 10^−5^) were downloaded from the GWAS catalog (04/06/2021). SNPs (MAF ≥ 0.01) that are in linkage disequilibrium (LD; R^2^ ≥ 0.8) with the NAFLD-associated SNPs (hereafter ‘NAFLD LD SNPs’ set) were identified using R package *LDlinkR*(31) (version 1.1.2). LD was calculated using the 1000 genomes phase III populations (AFR, AMR, EAS, EUR, SAS). The NAFLD-associated and NAFLD LD SNPs sets were combined and analyzed using the LGRN to identify the regulatory effects of these SNPs within the liver. We compared the impacts of the NAFLD-associated (and their LD SNPs) within the LGRN with the regulatory interactions that occur in diseased (NAFLD) livers. To do this, we tested to see if the eQTLs we identified within the LGRN were also found within an eQTL dataset derived from a cohort of NAFLD patients(32).

### Functional annotation of eQTLs and eGenes

eQTL functional classes were determined using the R programming interface of HaploReg (*HaploR*-4.0.2 package, 25/08/2021). Functional class annotations were based on the location of the eQTL relative to the gene(s) in the local vicinity.

eQTLs were annotated as enhancers if they overlap with ENCODE CHIP-seq narrow peaks corresponding to H3K27ac and H3K4me1 marks in liver tissue and HepG2 cell lines. Bed files from the following encode accessions were used for this annotation (*i*.*e*. ENCFF193GDV, ENCFF749VEQ, ENCFF413EGR, ENCFF953NPP, ENCFF625TWQ, ENCFF872NIB, ENCFF805YRQ, ENCFF452XJY, ENCFF287VIA).

The R package *biomaRt* (version 2.48.2) was used to classify genes as “protein-coding” (*i*.*e*. genes that encode functional protein products) and others (*i*.*e*. all other genes that do not encode functional proteins including pseudogenes, miRNA, rRNA, scRNA, function not yet determined [TEC], or uncategorized) according to their Ensembl gene annotations (URL: “https://may2021.archive.ensembl.org“, dataset: “*hsapiens_gene_ensembl*”)

Pathway enrichment analysis was performed using the R package *gprofiler2* (version 0.2.0) to identify pathways within the Kyoto Encyclopedia of Genes and Genomes (KEGG) that were significantly (FDR<0.05) enriched for the NAFLD-associated genes and their PPIN partners.

## Data and code availability

Supplementary Data 1 is available in figshare with the doi: https://doi.org/10.17608/k6.auckland.17087033

Supplementary Data 2 is available in figshare with the doi: https://doi.org/10.17608/k6.auckland.17087108

Supplementary Data 3 is available in figshare with the doi: https://doi.org/10.17608/k6.auckland.17087114

Supplementary Data 4 is available in figshare with the doi: https://doi.org/10.17608/k6.auckland.17087123

Supplementary Data 5 is available in figshare with the doi: https://doi.org/10.17608/k6.auckland.17087144

CoDeS3D pipeline is available at https://github.com/Genome3d/codes3d-v2.

Data and the scripts used for visualization are available at https://github.com/Genome3d/NAFLD-multimorbidities. Python v3.6.9 was used for all the python scripts. R v4.1.0 and RStudio v1.4.1717 was used for data analyses.

## Results

### Overview of the LGRN

The LGRN consisted of 327,145 eQTLs and 8,097 genes with 468,560 distinct spatially constrained eQTL target gene interactions (FDR < 0.05; Supplementary data 1 table 1). The majority (∼76%) of eQTLs within the LGRN have a single target gene (Figure 1A). This is consistent with observations for unconstrained *cis*-eQTLs within the GTEx liver dataset (Supplementary figure 1A). Approximately 92% (N = 229,376) and 95% (N = 73,749) of constrained eQTLs targeting one or more genes, respectively, are involved in *cis*-acting regulatory interactions. By contrast, 7% (N = 20,200) and 0.2% (N = 209) of the constrained eQTLs targeting one or multiple genes, respectively, are involved in *trans*-acting regulatory interactions. (Supplementary Figure 1B). These results are consistent with the liver gene regulatory landscape being predominantly comprised of constrained *cis*-acting eQTLs. However, this finding does not imply that either group of eQTLs is biologically more important than the other.

**Figure 1.**
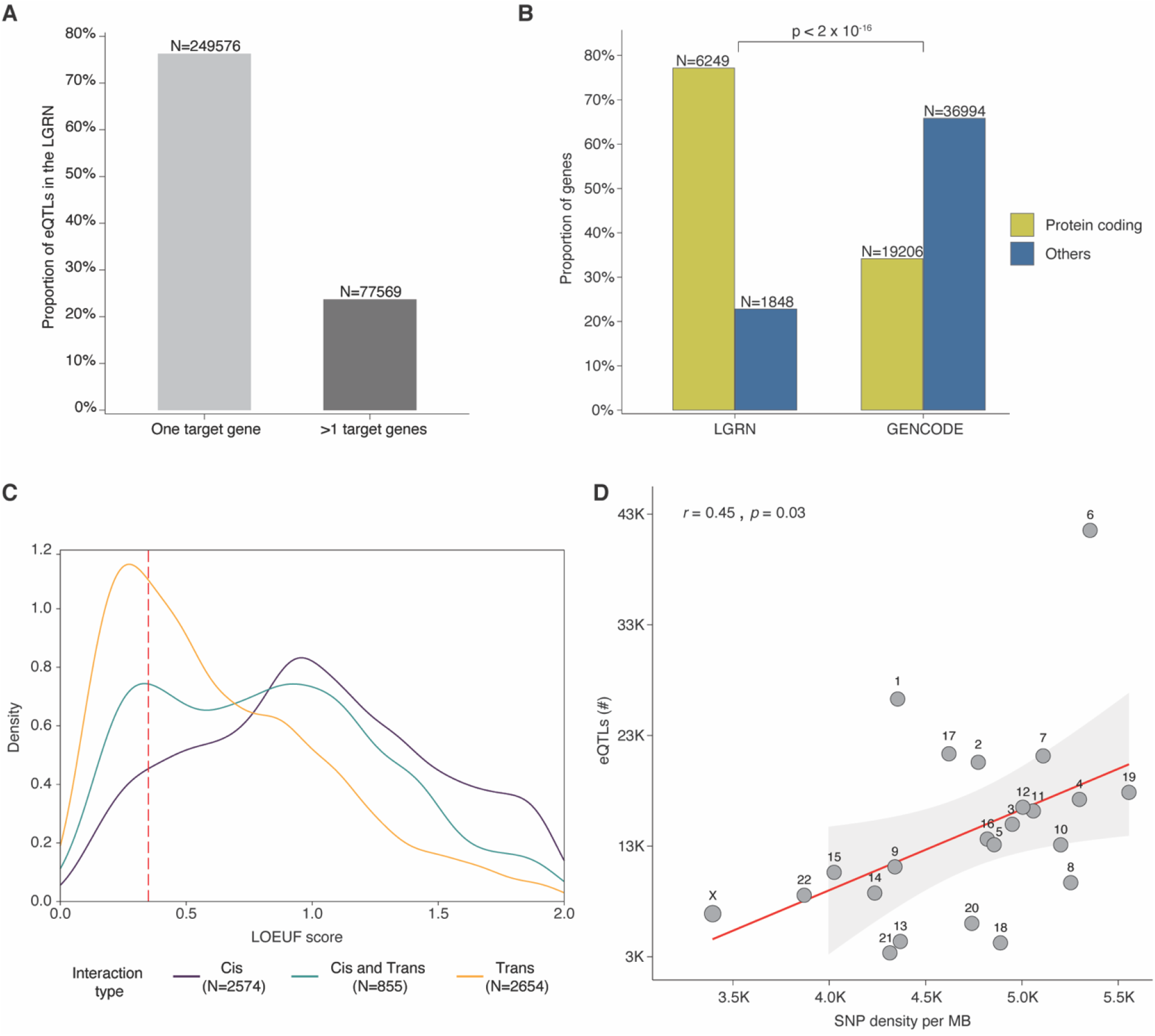
The eQTLs in the LGRN regulate the transcript levels of protein-coding genes that are evolutionarily constrained. **A**. The proportion of constrained eQTLs with only one target gene is higher than those with multiple target genes in the LGRN. **B**. Approximately 77% of the genes targeted by eQTLs in the LGRN are protein-coding, meaning that the LGRN eQTLs are significantly more likely to target protein-coding genes than those that do not encode proteins. **C**. Genes that are targeted by *trans*-acting eQTLs are highly intolerant to LoF mutations (LOEUF < 0.35) compared to the genes targeted by *cis*-acting and both *cis*-and *trans*-acting eQTLs. **D**. There is a positive correlative relationship between the number of spatially constrained eQTLs identified within the LGRN and the total number of common SNPs (MAF > 1%) on each chromosome, with several notable outliers (*e*.*g*. 1, 2, 6, 17 had more than expected. By contrast, 8, 13, 18, 20 and 21 had fewer than expected.).

Of the 8,097 genes in the LGRN, 50% (N = 4061) were altered by only *cis*-acting eQTLs, 13% (N = 1033) were affected by both *cis*- and *trans*-eQTLs and 37% (N = 3003) were affected by only *trans*-acing eQTLs (Supplementary Figure 1C). Within the LGRN, 77% of the eQTL target genes (N=6249) are protein coding (Figure 1B; Supplementary data 1 table 2). This is a significant bias towards protein-coding genes (two proportions *Z*-test, p < 2×10^−16^; Figure 1B). Remarkably, 90.38% of the target genes (N=2714 of 3003) of *trans*-acting eQTLs are protein-coding and 30.51% of these (N=828 of 2714) were intolerant to LoF mutations (Figure 1C; Supplementary figure 1D; Supplementary data 1 table 3). By comparison, only 65.43% (N = 2657 of 4061) of the *cis*-acting eQTLs target genes that encode proteins and only 9.52% of them (N = 253 of 2657) were intolerant to LoF mutations (Figure 1C; Supplementary figure 1D; Supplementary data 1 table 3). Therefore, in comparison with spatially constrained *cis*-eQTLs, spatially constrained *trans*-eQTLs predominantly target protein-coding genes and have a selection bias towards evolutionarily constrained genes.

We hypothesized that the number of liver-specific constrained-eQTLs identified on each chromosome would correlate with the number of common SNPs present on each chromosome. We observed a trend towards a positive correlation (Pearson’s *r*, 0.45, *p-*value = 0.03) between the number of SNPs and constrained eQTLs by chromosome. Notably, chromosomes 1, 2, 6 and 17 had significantly more eQTLs than expected, by contrast, chromosomes 8, 13, 18, 20 and 21 had fewer eQTLs (Figure 1D). It is interesting to speculate on the biological relevance of the increased numbers of constrained eQTLs within those chromosomes (*i*.*e*. 1, 2, 6 and 17), where there is a clear deviation from the chromosomes with similar number of SNPs. Similarly, we also observed a positive correlation (Pearson’s *r*, 0.7, *p-*value = 0.0002) between the number of genes and constrained eQTLs by chromosome (Supplementary figure 1E). Therefore, it remains possible that the increased numbers of eQTLs reflect gene density or heightened levels of transcriptional activity within these chromosomes.

### The regulatory impacts of NAFLD-associated SNPs differ with liver pathophysiological condition

The gene regulatory interactions for 739 SNPs (NAFLD-associated SNPs + NAFLD LD SNPs; collectively referred to as ‘NAFLD-SNPs’) (Supplementary data 2 table 1) were extracted from the LGRN (which was derived from a cohort of NAFLD unaffected individuals) and an eQTL-gene network derived from liver samples from a cohort of NAFLD patients(32) (hereafter “diseased-LGRN”; Supplementary data 2 table 2 and 3). Approximately 2% (N = 16 out of 739) of the NAFLD-SNPs were identified as eQTLs within the unaffected-LGRN (Figure 2A). By contrast, a significantly greater proportion (two proportion *Z*-rest, *p* = 1.192×10^−12^) of approximately 11% (N = 85 out of 739) were eQTLs within the diseased-LGRN (Figure 2B). The majority of these eQTLs (N = 11 [69%], unaffected-LGRN; N =80 [94%], diseased-LGRN) were located within intergenic regions of the genome (Supplementary figure 2A, B). Three of the unaffected- and 23 of the diseased-LGRN eQTLs were located within enhancer regions marked by H3K27ac and H3K4me1 modifications within HepG2 or liver cells (Supplementary figure 2C, D).

**Figure 2.**
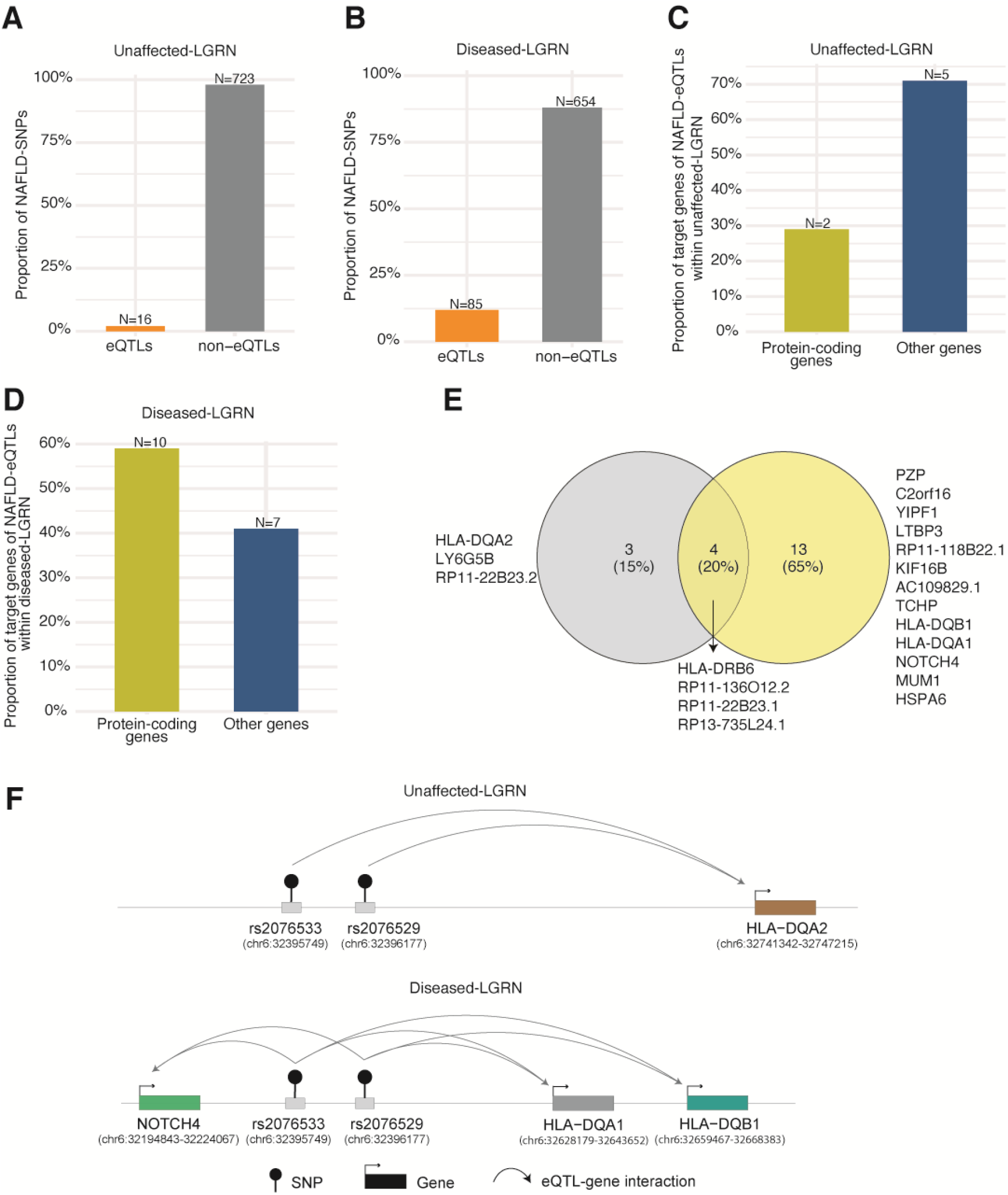
NAFLD-eQTLs exhibit context-specific regulatory effects. **A**. Approximately 2% of NAFLD-SNPs had gene regulatory effects within the unaffected-LGRN, and **B**. Approximately 11% of NAFLD-SNPs had regulatory effects on genes within the diseased-LGRN. However, in both datasets (**A** and **B**), significantly greater proportions (two proportions *Z*-test, *p* < 2.2×10^−16^) of NAFLD-SNPs were not associated with gene transcript levels. In both the **C**. unaffected-, and **D**. diseased-LGRN, NAFLD-eQTLs associate with transcript levels from both protein-coding and non-coding genes (*e*.*g*. non-coding RNAs, pseudogenes). **E**. Consistent with the context-specificity of eQTL effects, NAFLD-eQTLs had distinct sets of target genes in the unaffected- and diseased-LGRNs. **F**. NAFLD eQTL-gene regulatory interactions are context specific. The eQTLs that associated with *HLA-DQA2* transcript levels in the unaffected-LGRN were found to associate with *HLA-DQA1, -DQB1* and *NOTCH4* gene transcript levels within the diseased-LGRN. In this cartoon, the SNPs and genes are positioned according to relative order on chromosome 6.

The NAFLD-eQTLs from the unaffected- and diseased-LGRNs affect both protein-coding and other genes (*e*.*g*. non-coding RNAs, pseudogenes; Figure 2C, D). In total, 20 distinct genes were identified as being altered by NAFLD-eQTLs within the unaffected or diseased-LGRN. Amongst those genes, 13 and 3 were specific to the diseased- and unaffected-LGRNs respectively, and 4 genes were shared by both LGRNs (Figure 2E).

Amongst the genes that were affected by NAFLD-eQTLs in both the diseased and unaffected LGRNs, the transcript levels for *HLA-DRB6, RP11-22B23*.*1*, and *RP13-735L24*.*1* were associated with the same eQTLs (Supplementary data 2 table 2, 3). By contrast, *RP11-136O12*.*2* was regulated by distinct eQTLs (rs2980888, rs2954038 in the unaffected-LGRN; and rs2001845, rs2954018, rs2954026, rs2954032, rs2980867, rs55748921, rs56116731, rs7846466 in the diseased-LGRN). However, despite their being unique, the eQTLs that regulate *RP11-136O12*.*2* are located within a 31 kB block on chromosome 8 and are in strong LD (*R*^*2*^ > 0.8; calculated across AFR, AMR, EAS, EUR, SAS populations). Alternatively, the eQTLs (*e*.*g*. rs2076533, rs2076529) that associate with the transcript levels of *HLA-DQA2* gene, within the unaffected-LGRN, were associated with the transcript levels of different genes (*i*.*e. HLA-DQA1, HLA-DQB1* and *NOTCH4*) in the diseased-LGRN (Figure 2F). Collectively, these results suggest that the NAFLD-eQTLs’ gene regulatory effects are disease stage/context specific, reinforcing the need for greater longitudinal sampling during NAFLD development.

### NAFLD-candidate genes were targeted by eQTLs associated with traits/diseases that are linked to NAFLD

The LGRN was used to identify regulatory interactions (eQTLs) involving NAFLD-candidate genes (Table 1). Of 31 NAFLD-candidate genes, 13 (∼41%) were affected by spatially constrained eQTLs in the liver with 122 distinct eQTL-gene connections (Figure 3A and Supplementary data 3 table 1). Three genes (*i*.*e. IL32, TNFRSF12A, MBOAT7*) were impacted by *cis*-eQTLs, two (*i*.*e. EPB41L4A, THY1*) by both *cis* and *trans* eQTLs, and eight (*i*.*e. DTNA, COL1A1, COL1A2, PLPP4, CLIC6, SCTR, TM6SF2, PNPLA3*) by *trans*-eQTLs alone. Bootstrapping demonstrated that there is a negligible chance (bootstrapping; *p* = 0.0002) of finding 13 genes with eQTL connections in the LGRN for a set of 31 genes randomly selected from all genes [GENCODE release version 26; Figure 3B). Five of the candidate genes (*i*.*e. IL32, MBOAT7, EPB41L4A, THY1, HSD17B13*) were significantly associated with spatially unconstrained eQTLs within the liver regulatory map from GTEx (GTEx *q*-value threshold < 0.05; Supplementary figure 3A). However, this was not significantly different from what was expected by chance (bootstrapping; *p* = 0.0937; Supplementary figure 3B). Because of the requirement for spatially constrained eQTLs, some of the *cis*-eQTL-gene associations found in the GTEx liver regulatory map were not identified within the LGRN. For example, *HSD17B13* did not have any spatially constrained *cis*-eQTLs within the LGRN despite having significant unconstrained *cis*-eQTLs within the GTEx liver regulatory map. By contrast, *TNFRSF12A*, was found to be associated with spatially constrained *cis*-eQTLs within the LGRN (FDR < 0.05) despite having no significant *cis*-eQTLs contacts within the GTEx liver regulatory map (GTEx *q*-value threshold > 0.05). Overall, analysis of the *cis* and *trans*-acting constrained eQTLs within the LGRN enabled the identification of putative regulatory sites for the NAFLD-candidate genes that were not captured by classical eQTL analyses.

**Figure 3.**
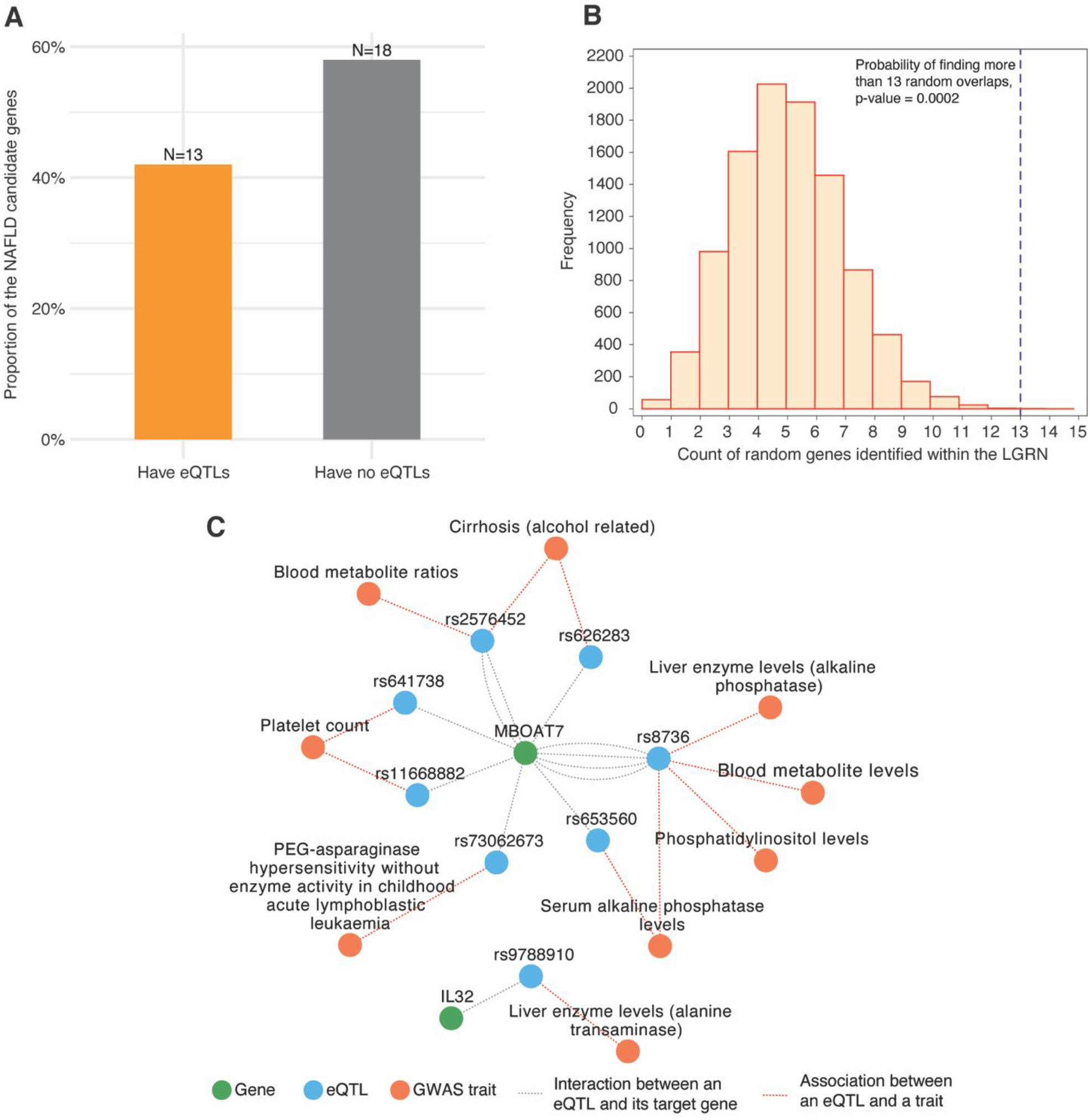
The constrained eQTLs regulating the NAFLD-candidate genes within the LGRN were enriched for traits closely linked to NAFLD. **A**. The transcript levels of 13 of the 31 NAFLD-candidate genes were altered by constrained eQTLs within the LGRN. **B**. Bootstrapping (N = 10000) demonstrated that the identification of constrained eQTLs for 13 NAFLD-candidate genes within the LGRN was significantly greater than expected by chance. The number of gene overlaps detected between each random gene set and the genes within the LGRN (identified by the bootstrap resampling method) is distributed around five (*mean* = 4.614), which is lesser than that of the observed overlaps (N = 13; dotted line). This means that the chance of identifying the 13 genes (from a set of 31 random genes) with eQTL connections within the LGRN is low (*p* = 0.0002). **C**. The constrained eQTLs (blue) in the LGRN that affect the transcript levels of *MBOAT7* and *IL32* (green) were significantly (*p* < 0.026) associated with traits (orange) that are closely linked to NAFLD. Significance of the enrichment of eQTLs within trait associated SNP sets was calculated using a hypergeometric test followed by Benjamini-Hochberg FDR correction.

We investigated whether the 122 eQTLs that were associated with the transcript levels of the 13 NAFLD-candidate genes had been previously associated with NAFLD or other complex traits by GWAS. We performed a hypergeometric test to identify enrichment of the NAFLD-candidate gene eQTL associated traits amongst all traits within the GWAS catalog. The enrichment results showed that none of the 122 eQTLs was associated with, or showed enrichment for, NAFLD as a trait within the GWAS catalog (Supplementary data 3 table 2). Moreover, although genes (*e*.*g. PNPLA3, TM6SF2, GCKR, HSD17B13*) have been mapped to NAFLD-associated variants by GWAS, these mappings are not supported in the LGRN. However, eQTLs associated with the changes in *MBOAT7* and *IL32* transcript levels were enriched (FDR corrected hypergeometric test, *p* < 0.026) for associations with liver enzyme levels (*i*.*e*. alanine transaminase and alkaline phosphatase), platelet count and blood metabolite levels (Figure 3C, Supplementary data 3 table 2). The eQTLs for *MBOAT7* (rs641738, associated with platelet count; and rs2576452, associated with blood metabolite ratios), and *IL32* (rs9788910, associated with liver enzyme levels) are in an H3K4me1 modified enhancer regions in liver-derived cells. Collectively, these results are consistent with the hypothesis that the eQTLs of *MBOAT7* and *IL32* affect liver function and contribute to the multimorbid relationship of these traits to NAFLD.

### PPIN partners of the NAFLD-candidate genes are targeted by eQTLs enriched for the clinical traits linked to NAFLD

We hypothesised that additional modifiers of NAFLD would be identified by looking at the functional partners of the 13 NAFLD-candidate genes (controlled by constrained eQTLs) within a liver-specific PPIN. One hundred and eleven (111) genes were identified, within the liver-specific PPIN, as encoding proteins that directly interact with 11 of the NAFLD-candidate genes (*i*.*e. TNFRSF12A, IL32, SCTR, PNPLA3, COL1A1, MBOAT7, EPB41L4A, DTNA, THY1, CLIC6, COL1A2*; Supplementary data 4 table 1). Functional analysis of the 122 genes (*i*.*e*. 111 interacting genes and the 11 NAFLD-candidate genes) from the liver PPIN identified enrichment (FDR corrected, *p* < 0.05) within molecular pathways affecting liver function (*e*.*g*. focal adhesion, AGE-RAGE signaling pathway in diabetic complications, platelet activation) and other pathways linked to NAFLD pathogenesis (*e*.*g*. PI3K-Akt signaling pathway, phospholipase D signaling pathway; Figure 4A and Supplementary data 4 table 2). Among the 36 genes overrepresented in the pathways only two (*COL1A1, COL1A2*) were NAFLD-candidate genes. Notably, *DTNA, TNFRSF12A* and *COL1A1* interacted with the genes (*e*.*g. BRCA1, TRAF1, TRAF2, STAT3*) that were enriched within cancer-related pathways (*e*.*g*. PI3K-Akt signaling pathway, transcriptional misregulation in cancer, proteoglycans in cancer; Supplementary data 4 table 1, 2).

**Figure 4.**
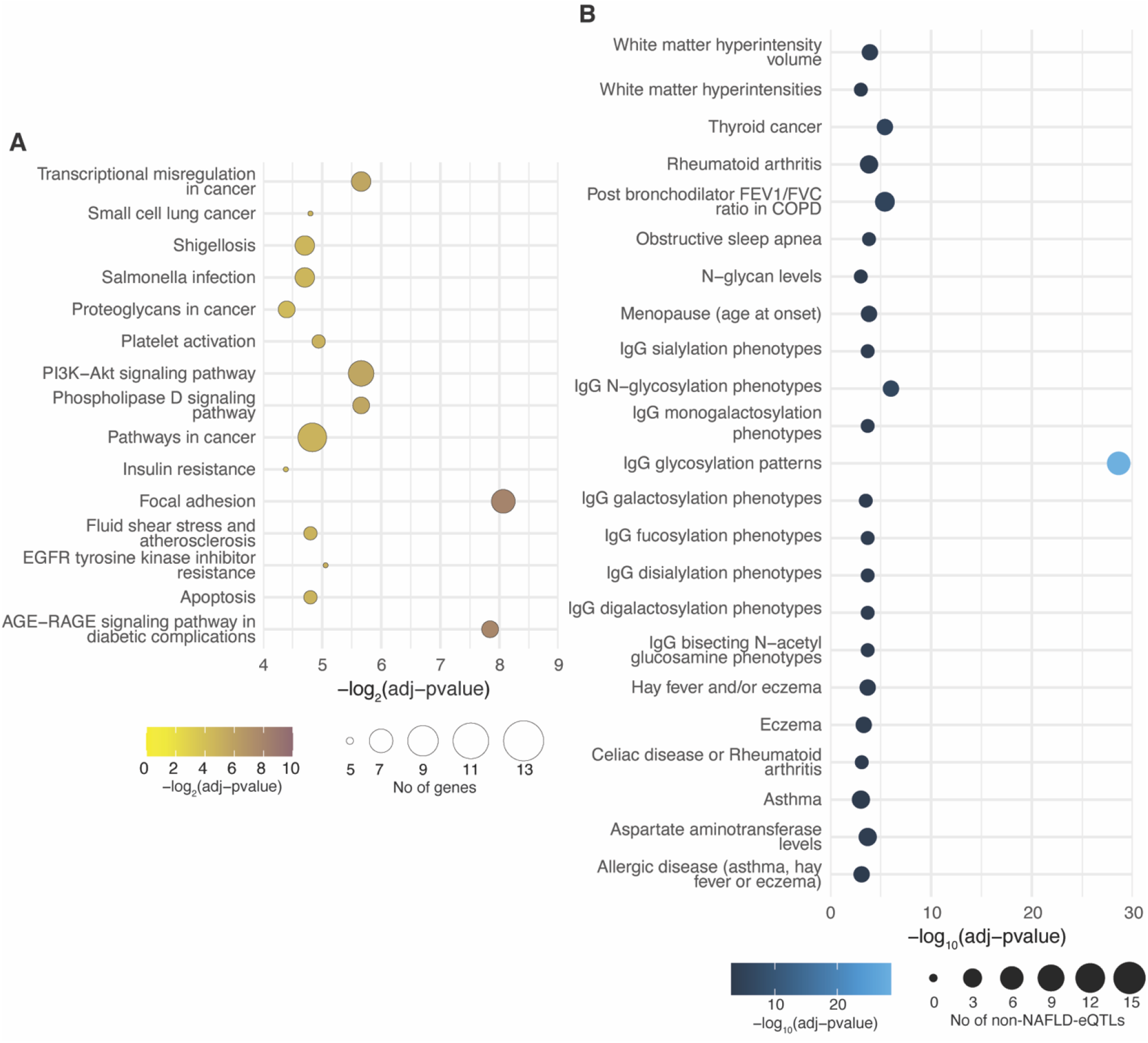
Analysis of the PPIN partners of NAFLD-candidate genes revealed genetic links between NAFLD and clinical traits relevant to NAFLD. **A**. NAFLD-candidate genes and their PPIN partners are enriched (FDR corrected, *p* < 0.05) for the KEGG biological pathways critical in liver function and NAFLD pathogenesis (Supplementary data 4 table 2). **B**. The constrained eQTLs controlling the expression of the PPIN partner genes of the NAFLD-candidate genes were significantly enriched (FDR corrected hypergeometric test, *p* < 0.05) for clinical traits that have been associated with NAFLD (Supplementary data 4 table 3).

We assessed the traits associated with constrained eQTLs within the LGRN that altered transcript levels of the interacting partners of the NAFLD-candidate genes. The eQTLs affecting the interacting partners were significantly enriched (FDR corrected hypergeometric test, *p* < 0.05) for variants associated by GWAS with traits including IgG glycosylation patterns, IgG N-glycosylation phenotypes, N-glycan levels, OSA, aspartate aminotransferase levels (Figure 4B; Supplementary data 4 table 3). These traits have previously been linked to NAFLD(10,33–35). The eQTLs were also significantly enriched for variants associated with allergic and autoimmune diseases, thyroid cancer, and white matter lesions (Figure 4B and Supplementary data 4 table 3). Notably, these conditions have been epidemiologically or mechanistically linked to metabolic syndrome(36– 38), one of the major NAFLD risk factors. Collectively, these results are consistent with the overall risk or severity of NAFLD arising from the convergence of NAFLD related traits and candidate genes through shared impacts on PPINs that are enriched within biological pathways affecting liver functions.

### Shared gene regulatory mechanisms link NAFLD and other related complex traits

We hypothesized that additional eQTLs associated with the expression of NAFLD eQTL target genes would modify NAFLD complications. To test this hypothesis, we interrogated the unaffected-LGRN to identify the eQTLs associated with the transcript levels of the NAFLD-eQTL target genes (excluding the NAFLD eQTLs themselves). We then mined the GWAS catalog to identify the traits and conditions that were enriched for these eQTLs. The *HLA* genes were excluded from this analysis to minimize bias towards overstudied traits/diseases. Of the 16 genes we analysed, the transcript levels of 11 (*LY6G5B, RP11-22B23*.*2, RP11-136O12*.*2, RP11-22B23*.*1, RP13-735L24*.*1, C2orf16, NOTCH4, PZP, MUM1, TCHP, KIF16B*) were associated with 674 non-NAFLD-eQTLs (Supplementary data 5 table 1). Trait enrichment analysis identified significant enrichment for GWAS traits that are relevant to NAFLD (*e*.*g*., plasma omega-6 polyunsaturated fatty acid levels (adrenic acid), phospholipids levels, obesity (extreme), coronary artery disease, body fat percentage) for 26 of the 37 non-NAFLD-associated eQTLs that affect transcript levels of *C2orf16, LY6G5B, NOTCH4, RP11-22B23*.*1, RP11-22B23*.*2, RP13-735L24*.*1* (Figure 5 and Supplementary data 5 table 2). eQTLs (N = 8) targeting three non-coding genes, *RP11-22B23*.*1, RP11-22B23*.*2, RP13-735L24*.*1* were significantly enriched (FDR corrected hypergeometric test, *p*-value = 5.828×10^−18^) for time-dependent creatinine clearance change response to tenofovir treatment in HIV infection (Supplementary data 5 table 2). Our discovery-based approach also identified enrichment for traits that have not yet been linked to NAFLD clinically or epidemiologically (*e*.*g*. myopia) (Figure 5). Collectively, these results revealed underlying shared gene regulatory mechanisms that link NAFLD with known and previously unidentified comorbidities or complications.

**Figure 5.**
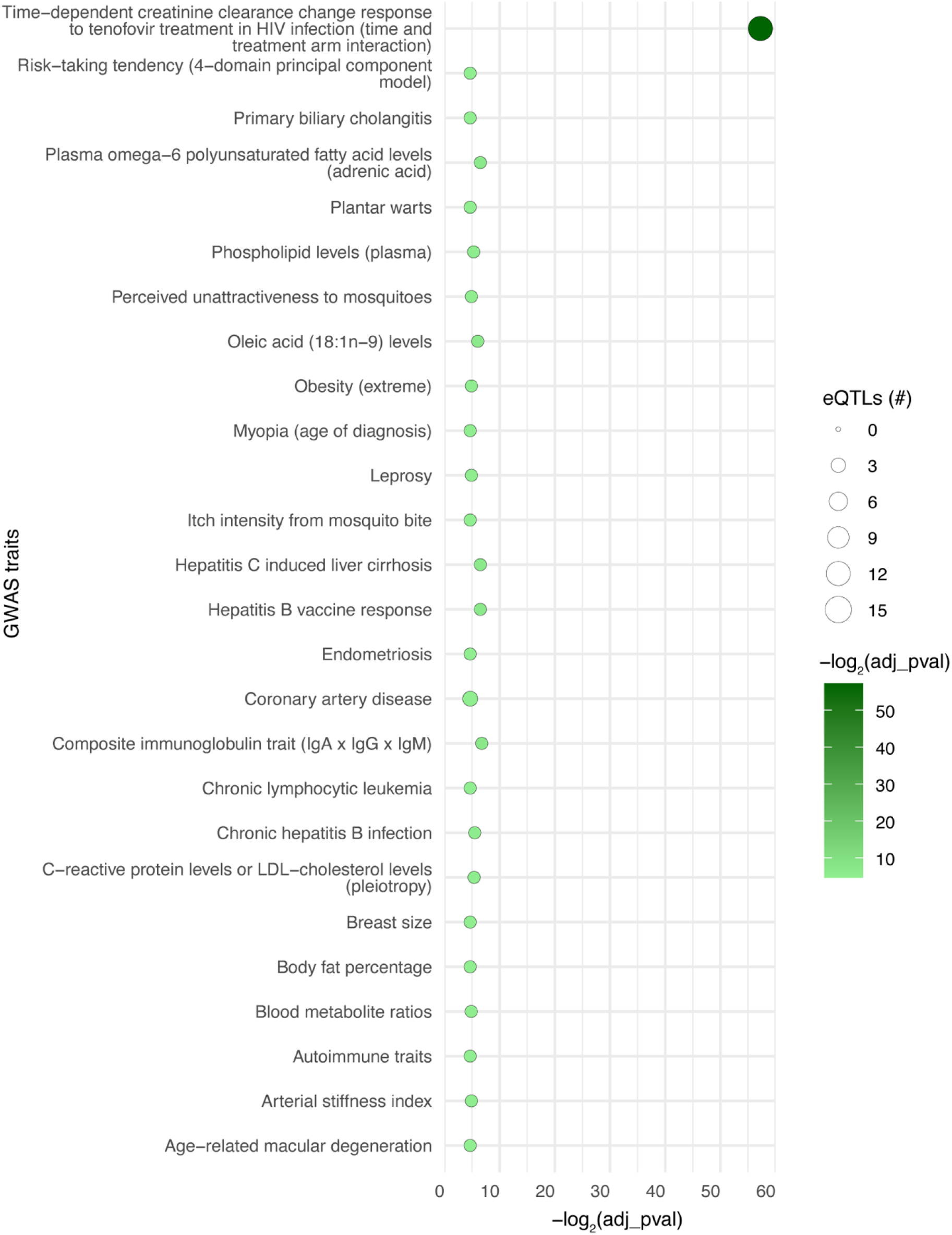
NAFLD- and comorbid trait-associated eQTLs share common gene targets within the unaffected-LGRN. Transcript levels of the genes affected by NAFLD-eQTLs within unaffected- and diseased-LGRN were also altered by eQTLs that are significantly (*p* < 0.05) enriched for traits linked to NAFLD. Hypergeometric test followed by multiple correction was performed to determine the traits overrepresented by non-NAFLD-eQTLs that altered the expression levels of NAFLD-eQTL target genes within unaffected-LGRN.

## Discussion

We used constrained eQTLs to establish a LGRN from individuals without NAFLD. The LGRN was then used to identify regulatory connections involving NAFLD-candidate genes. Comparisons of LGRNs from unaffected- and diseased-individuals demonstrated that the NAFLD-SNPs establish distinct gene regulatory interactions that are dependent upon disease state. The NAFLD-candidate genes and their interacting partners within the liver-specific PPIN were regulated by eQTLs that were enriched for associations with phenotypes relevant in NAFLD, but not NAFLD itself. Finally, we demonstrated that the genes dysregulated by NAFLD-eQTLs were affected by additional eQTLs that were associated with traits/diseases related to NAFLD. Overall, our approach enabled the *de novo* identification of the complex gene regulatory mechanisms linking NAFLD with comorbid conditions through genetic and biological pathways.

Within the LGRN, NAFLD-candidate genes and the proteins they interact with were dysregulated by eQTLs that were enriched for traits/diseases associated with NAFLD, but not NAFLD itself. For example, constrained eQTLs affecting the transcript levels of genes *MBOAT7* and *IL32* were significantly enriched within SNPs associated with phosphatidylinositol levels, serum alkaline phosphatase levels, platelet count, and blood metabolite levels. These traits are known to negatively affect liver functions(39–41). Given that *MBOAT7* and *IL32* genes play critical roles in the pathogenesis and progression of NAFLD(42,43), the regulatory interactions captured between eQTLs associated with liver transformations and these genes provide objective explanations as to how these traits might genetically influence an individual’s susceptibility to NAFLD.

Previous studies have shown that NAFLD is associated with an increased risk of extrahepatic cancers(44). Consistent with this, we show that the gene products of NAFLD-candidate genes interact with proteins that are enriched for cancer-related pathways (*e*.*g*. pathways in cancer [KEGG: hsa05200], Small cell lung cancer [KEGG: hsa05222]) within the liver-specific PPIN. Given that proteins that physically interact within the PPIN tend to be involved in the same cellular process(45,46), NAFLD-candidate genes may function within cellular pathways involved in cancers.

We did not identify any regulatory interactions between NAFLD-SNPs and NAFLD-candidate genes within the unaffected- or diseased-LGRNs. Therefore, the risk that is attributable to the NAFLD-associated SNPs appears to be mechanistically independent of the NAFLD-candidate genes, within the gene regulatory networks from both diseased and unaffected individuals. NAFLD-eQTLs altered *HLA* genes (*HLA-DQA2, HLA-DQA1, HLA-DQB1, HLA-DRB6*) within both diseased and unaffected individuals. Despite this, the majority of the eQTLs identified in diseased-LGRN were not associated with gene expression changes in the unaffected-LGRN. For example, rs965813 is significantly associated with changes in expression of *C2orf16* only in patients with NAFLD. This highlights the fact that certain eQTLs are active only in the context of disease(47) and highlights the involvement of immune processes in the risk and pathogenesis of NAFLD.

NAFLD-eQTL target genes were also regulated by additional eQTLs that were significantly enriched within SNP sets for traits or diseases related to NAFLD pathology (*e*.*g*. extreme obesity, LDL-cholesterol levels, coronary artery disease). This suggests that the eQTLs associated with NAFLD and conditions linked to it may initiate phenotypic manifestations by acting on shared gene targets. Whilst intriguing, the identification of putative eQTL effects does not prove causation. Therefore, further experimental investigations are required to establish causal relationships. This is particularly important as our discovery-based approach has identified novel associations that are seemingly unrelated to NAFLD (*e*.*g*. breast size), but may provide important insights into this complex disorder.

Our approach is not without limitations. Principle amongst them was the assumption that there was a physical interaction captured between variants and target gene. Therefore, the LGRN we constructed does not contain variant-gene pairs that do not require a physical interaction. As such, we may have missed loci that influence their target genes through alternative mechanisms such as alternative splicing, alternative polyadenylation(48,49). Notwithstanding this limitation, we contend that the LGRN we constructed explains a substantial proportion of the impact of common SNPs (MAF>0.01) on gene regulation in the liver. Additionally, we have tried to mitigate this limitation by comparing our results directly with the GTEx consortium findings to provide context and highlight the overlap with the traditional (unconstrained) eQTL analysis. Secondly, our assumption that eQTL associated changes in gene transcript levels correlate with changes in protein levels may not always hold true. Gene expression is the result of a series of connected biological processes (*e*.*g*. transcription, mRNA degradation etc.) each of which can dissociate protein and transcript abundance(50,51). Thirdly, we lacked a liver tissue-specific eQTL dataset from individuals who were screened for NAFLD and confirmed as being non-NAFLD control. Instead, we used the GTEx liver eQTL map as a representative network for the NAFLD-unaffected group. Lastly, the differences we observed in the regulatory effects of the same NAFLD-eQTLs in the unaffected- and diseased-LGRNs may be attributable to differences between the study populations. Specifically, the unaffected-LGRN was derived from a cohort of predominantly European ancestry(25). By contrast, the diseased-LGRN is derived from a predominantly East Asian cohort(32). It is possible that this could have resulted in the identification of distinct eQTLs (within the unaffected- and diseased-LGRNs) that were in strong LD (*R*^*2*^ > 0.8), affecting the same gene. Despite these limitations, we identified statistically robust trait associations for the eQTLs that share gene regulatory mechanisms with NAFLD-eQTLs, as well as those that affect NAFLD-candidate genes (and their PPIN partners). This indicates the potential biological significance of the genetic links that were identified as being co- or multimorbid with NAFLD.

In conclusion, our approach systematically integrated three-dimensional chromatin contact, eQTL and protein interaction network data into a LGRN to uncover traits that are genetically linked with NAFLD. This enabled the *de novo* identification of comorbid traits, some of which are known and others of which are novel. Together, this approach provides novel insights into NAFLD and its multimorbid conditions.

## Supplementary figures

**Supplementary figure 1.**
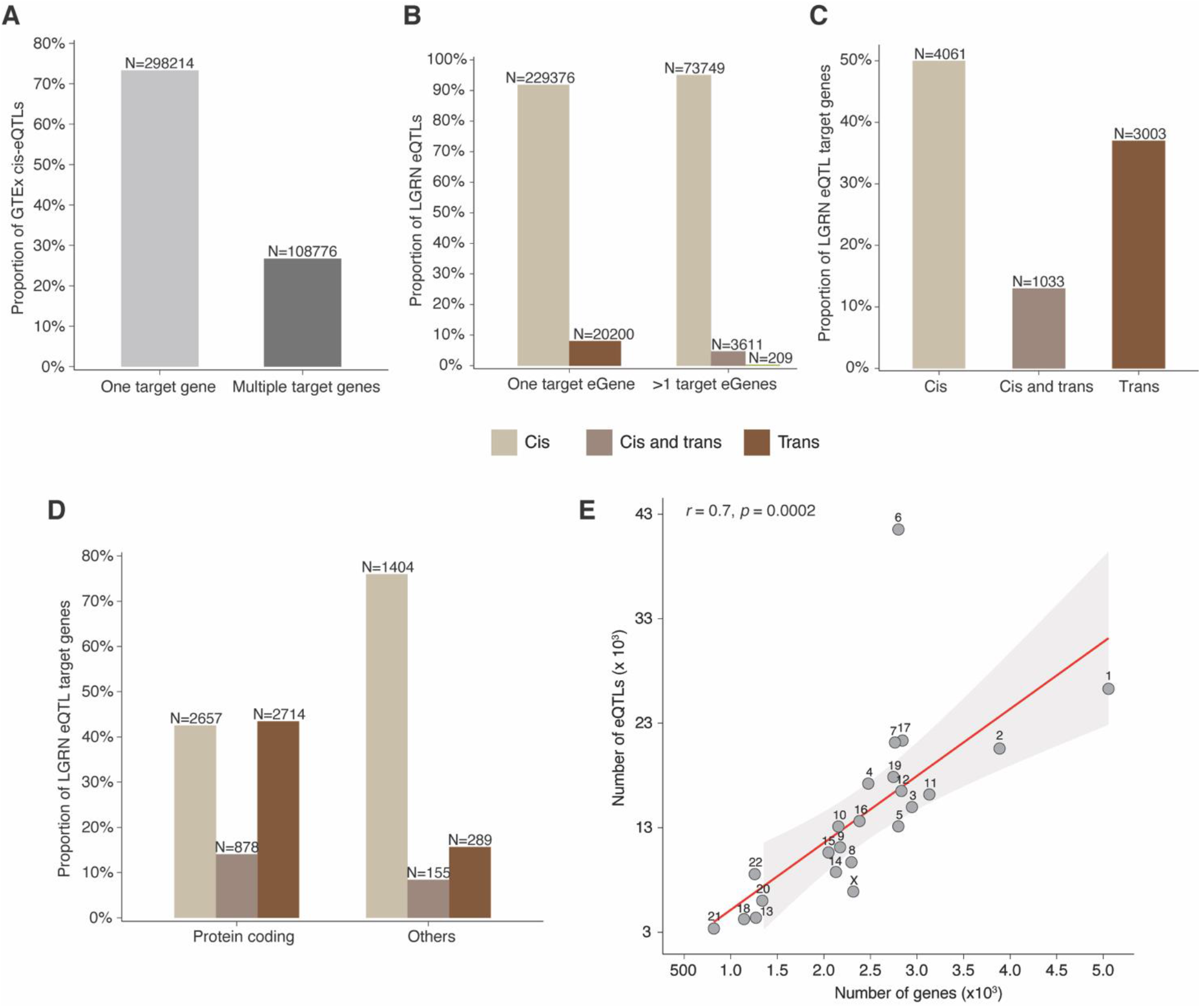
Both *cis* and *trans*-acting eQTLs contribute to the dysregulation of protein-coding genes in the liver. **A**. The proportion of cis-eQTLs, identified by the classical linear regression method within the GTEx liver samples (V8), that have only one or multiple target genes is equivalent to that observed for the LGRN. **B**. In comparison to *trans*-eQTLs, a large proportion of the *cis*-eQTLs are associated with the expression levels of either single or multiple target genes in the LGRN, consistent with gene regulatory connections within the LGRN being predominantly controlled by *cis*-acting eQTLs. **C**. The majority of genes within the LGRN are only targeted by *cis*-eQTLs. However, a large number of genes are also affected by *trans*-eQTLs, suggesting a role for long-range regulation in controlling gene expression within the liver. **D**. *Trans*-acting eQTLs, despite being modest in quantity within the LGRN, affect a significantly greater proportion (two-sample proportion *Z-*test, *p* = 1.45 ×10^−130^) of protein-coding genes than the cis-eQTLs. **E**. There is a positive correlation between the number of spatially constrained eQTLs identified within the LGRN and the total number of genes (according to GENCODE V26) on each chromosome. Again chromosomes 6 and 17 have significantly more eQTLs than expected.

**Supplementary Figure 2.**
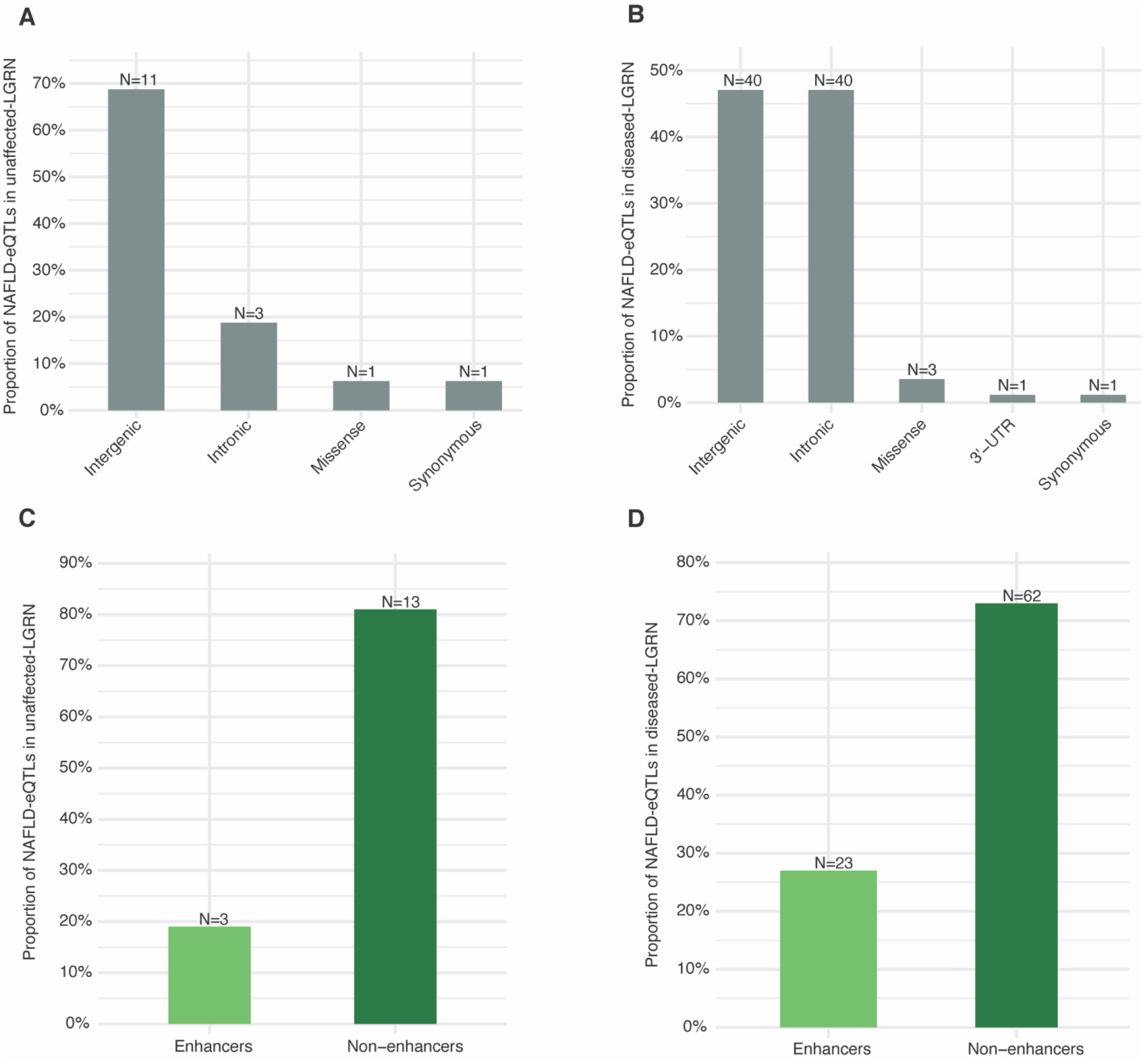
NAFLD-eQTLs are located in regulatory regions within the genome. NAFLD-eQTLs within the **A**. unaffected- and **B**. diseased-LGRNs were primarily located in the intergenic (*i*.*e*. the non-coding segment of the genome between two genes), intronic or intragenic (*i*.*e*. non-coding segments of a gene) regions of the genome. Approximately, 19% and 27% of the NAFLD-associated eQTLs within the **C**. unaffected- and **D**. diseased-LGRNs, respectively, reside in genomic regions tagged by H3K27ac and H3K4me1 (e.g. “enhancers”) in the liver cells, consistent with the putative gene regulatory effects of NAFLD-eQTLs in the hepatic cells.

**Supplementary figure 3.**
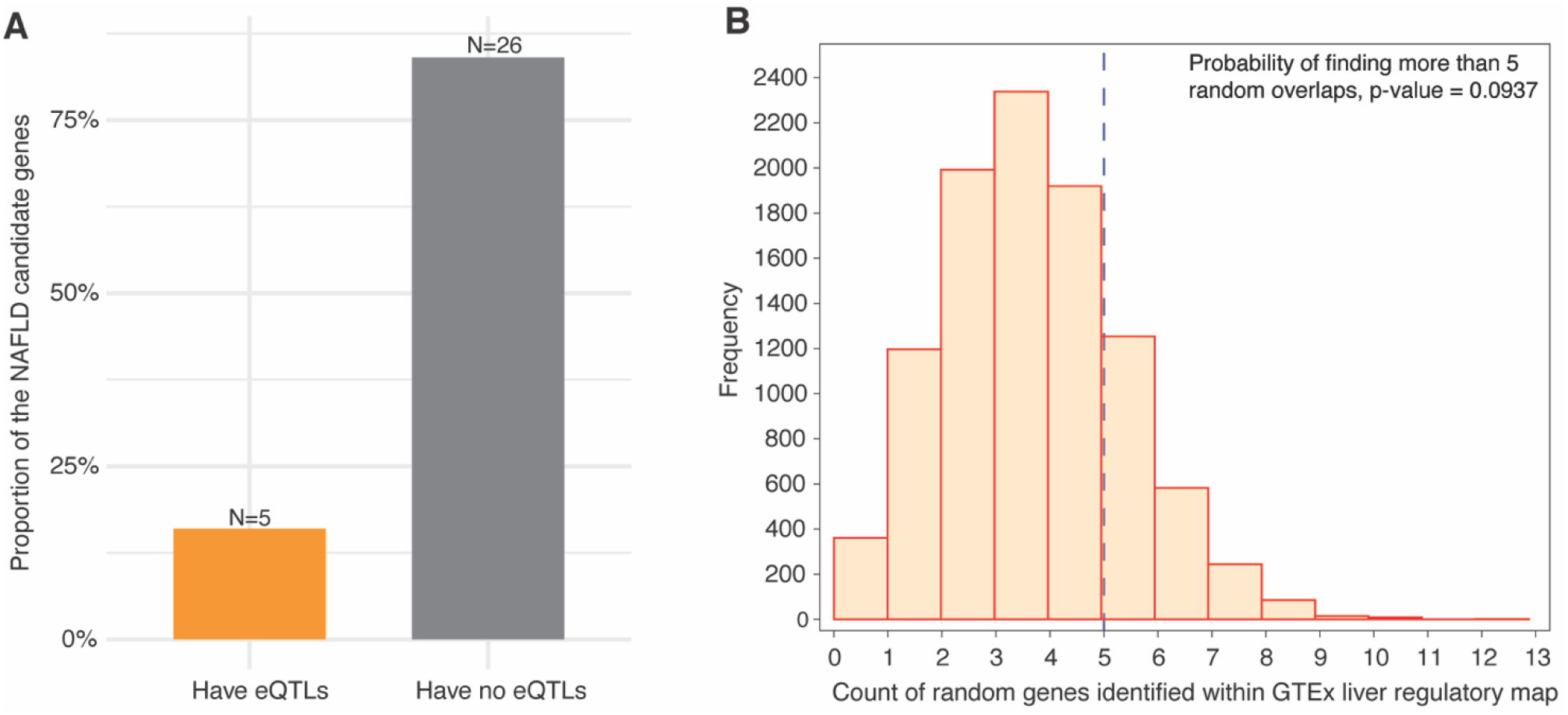
There is no significant enrichment for NAFLD-candidate genes targeted by spatially unconstrained *cis*-eQTLs within the GTEx liver map. **A**. Five of the 31 NAFLD-candidate genes expression levels were altered by spatially unconstrained *cis*-eQTLs within the GTEx liver map. **B**. Bootstrapping demonstrated that the identification of unconstrained *cis*-eQTLs for NAFLD-candidate genes within the GTEx liver map was not greater than expected by chance. Bootstrapping identified that the number of gene overlaps detected between each random gene set and the genes within the GTEx liver regulatory map is distributed around four (*mean* = 6.723). Therefore, the identification of five genes (from a set of 31 random genes) with eQTL connections within the GTEx liver regulatory map is not statistically significant (*p* = 0.0937). Significance of the enrichment for traits was calculated using a hypergeometric test followed by Benjamini-Hochberg FDR correction.

## Author Contributions

SG conceptualized, performed analyses, interpreted data, and wrote the manuscript. WS contributed to conceptualization, data interpretation and manuscript revision.TF and REZ contributed to statistical analysis and commented on the manuscript. MC contributed to data analysis and commented on the manuscript. JOS directed the study, contributed to data interpretation, and co-wrote the manuscript. All authors contributed to the article and approved the submitted version.

